# Construction of the third generation *Zea mays* haplotype map

**DOI:** 10.1101/026963

**Authors:** Robert Bukowski, Xiaosen Guo, Yanli Lu, Cheng Zou, Bing He, Zhengqin Rong, Bo Wang, Dawen Xu, Bicheng Yang, Chuanxiao Xie, Longjiang Fan, Shibin Gao, Xun Xu, Gengyun Zhang, Yingrui Li, Yinping Jiao, John Doebley, Jeffrey Ross-Ibarra, Vince Buffalo, M. Cinta Romay, Edward S. Buckler, Yunbi Xu, Jinsheng Lai, Doreen Ware, Qi Sun

## Abstract

**Background:** Characterization of genetic variations in maize has been challenging, mainly due to deterioration of collinearity between individual genomes in the species. An international consortium of maize research groups combined resources to develop the maize haplotype version 3 (HapMap 3), built from whole genome sequencing data from 1,218 maize lines, covering pre-domestication and domesticated Zea mays varieties across the world.

**Results:** A new computational pipeline was set up to process over 12 trillion bp of sequencing data, and a set of population genetics filters were applied to identify over 83 million variant sites.

**Conclusions:** We identified polymorphisms in regions where collinearity is largely preserved in the maize species. However, the fact that the B73 genome used as the reference only represents a fraction of all haplotypes is still an important limiting factor.

## BACKGROUND

Maize, one of the most important cereals in the world, also happens to be among the crop species with the most genetic diversity. Advances in the next generation sequencing technologies made it possible to characterize genetic variations in maize at genomic scale. The previously released maize HapMap2 were constructed with whole genome sequencing data of 104 maize lines across pre-domestication and domesticated *Zea mays* varieties [1]. Since then, more maize lines have been sequenced by the international research community, and a consortium was formed to develop the next generation haplotype map. The maize HapMap 3 consortium includes, among others, BGI-Shenzen, Chinese Academy of Agricultural Sciences, China Agricultural University, International Maize and Wheat Improvement Center (CIMMYT). High-coverage data for 31 European and US Flint and Dent lines is also available in Ref. [2]. Altogether, in this work we used a total of 1218 maize lines sequenced with depth varying from below 1x to 59x.

A common approach in today’s genetic diversity projects is to map the shotgun sequencing reads from each individual onto a common reference genome to identify DNA sequence variations, and the physical positions of the reference genome is used as a coordinate system for the polymorphic sites. A good example is the human 1000 genome project [3]. The computational data processing pipeline developed for the human project, GATK, has been widely adopted for identifying genetic variations in many other species [4].

As the sequencing technology is improved and sequencers’ base calling error model gets more accurate, the computational challenges in genotyping by short-read sequencing have shifted from modeling sequencer machine artifacts errors to resolving genotyping errors derived from incorrect mapping of short reads to the reference genome. The problem is associated with the experimental design that uses the single-reference genome as coordinate system. Taking maize as an example, the reference being used is a 2.1 Gb assembly from an elite inbred line B73 that represents 91% of the B73 genome [5], and was estimated to capture only ~70% of the low-copy gene fraction of all inbred lines [6]. The sequence alignment software, however, can map 95-98% of the whole genome sequencing reads to the reference. That suggests a high percentage of the reads were mapped incorrectly, either being mapped to the paralogous loci or highly repetitive regions under-represented in the reference assembly. The genetic variants called from the miss-mapped reads need to be corrected computationally. The maize HapMap2 relied on linkage disequilibrium in the population to purge most of the bad markers caused by alignment errors. To construct maize HapMap 3, a new computational pipeline was developed from scratch to handle the sequencing data from 10 times more lines, and also took advantage of the high quality genetic map constructed from the GBS technology [7, 8] which was not present when HapMap2 was constructed.

Genome structure variation in the population, including transposition, deletion, duplication and inversion of the genomic segments, poses another challenge in the HapMap projects. As the physical genomes of each of the individuals included in the HapMap projects vary both by size and structure, and there is no co-linearity of all the sequence variants between the reference and genomes of each of the individuals, it is not always possible to anchor all genetic variants in a population onto a single reference coordinate system. As a compromise, markers included in the maize HapMap are defined as sites of which the physical positions of the B73 alleles matching the markers’ consensus genetic mapped positions.

Here we present maize haplotype map version 3 (HapMap 3), which is a result of coordinated efforts of the international maize research community. The build includes 1,218 lines and over 83 million variant sites anchored to the B73 reference genome version AGP v3.

## DATA DESCRIPTION

The sequencing data used in this work is comprised of 12,497 billion base pairs in a total of 113,702 billion Illumina paired-end reads, originating from 1,218 maize and teosinte lines. The data was collected from several sources over several years, and varies in quality, read length, and coverage. Basic information about various datasets and stages of the HapMap 3 project they were used in are summarized in Table 1. Each of the 1,218 lines were sequenced at depth varying from below 1x to 59x, using reads of lengths ranging from 44 through 201bp, averaging 110 bp. All reads were aligned to maize reference genome B73 version AGP v3 using BWA mem aligner [9]. Overall, 95-98% of the reads were mapped to the reference genome, although only about 50-60% with non-zero mapping quality.

**Table 1:**
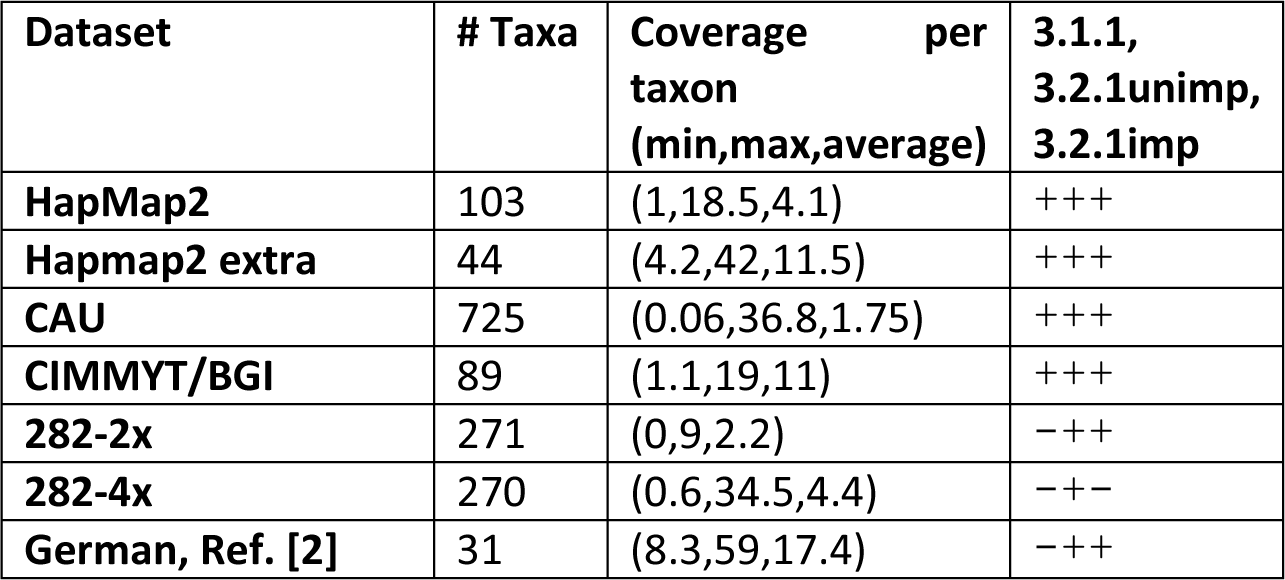
Sequence datasets used in various stages of HapMap 3

Taxa from sets “HapMap2”, “HapMap2 extra”, and “CAU” partially overlap. The “282” libraries, sequenced twice represent 271 taxa. A “+” means that the dataset was used in a given stage, “-“ that it was not.

## ANALYSIS

### Initial variant discovery

The HapMap 3 pipeline is summarized in Figure 1. First, polymorphic sites were called for a set of 916 taxa from datasets HapMap2 through CIMYYT/BGI (7,191 billion base pairs, 74,643 million reads). In the first step, a custom built software tool was used to determine genotypes for each taxon at each site of the genome based on allelic depths at that site. Bases counted towards depth had base quality score of at least 10 and originated from reads with mapping quality at or above 30. Only sites where at least 10 taxa had coverage of 1 or more were considered. The allelic depths were subject to segregation test (ST – see next section), which determines the probability that a given distribution of allelic depths over taxa has been obtained by chance. Sites with high probability, which are likely a result of random sequencing errors, have been eliminated by applying a p-value threshold of 0.01. In this first round, a total of 196 million tentative polymorphic sites were selected. In the second step, these sites were filtered using the identity by descent (IBD) information derived from about 0.5 million of high-quality polymorphisms obtained previously [8] using the Genotyping-By-Sequencing (GBS) approach [7]. These GBS variants (GBS anchor) were used to determine regions of IBD, where certain pairs of taxa are expected to have identical haplotypes. The raw tentative polymorphisms violating these IBD constraints were then filtered out, leaving 96.8 million sites. At roughly half of the sites surviving this filter, minor allele was not present in IBD contrasts. Such sites, typically with low minor allele frequency, are less reliable and have been marked with “IBD1” flag in the VCF files (see Table 2 for summary of flags and parameters present in HapMap 3 VCF files). The ST- and IBD-filtered variant sites were then used in two separate procedures, leading to two versions of HapMap 3 genotypes, referred to as HapMap 3.1.1 and HapMap 3.2.1.

**Figure 1:**
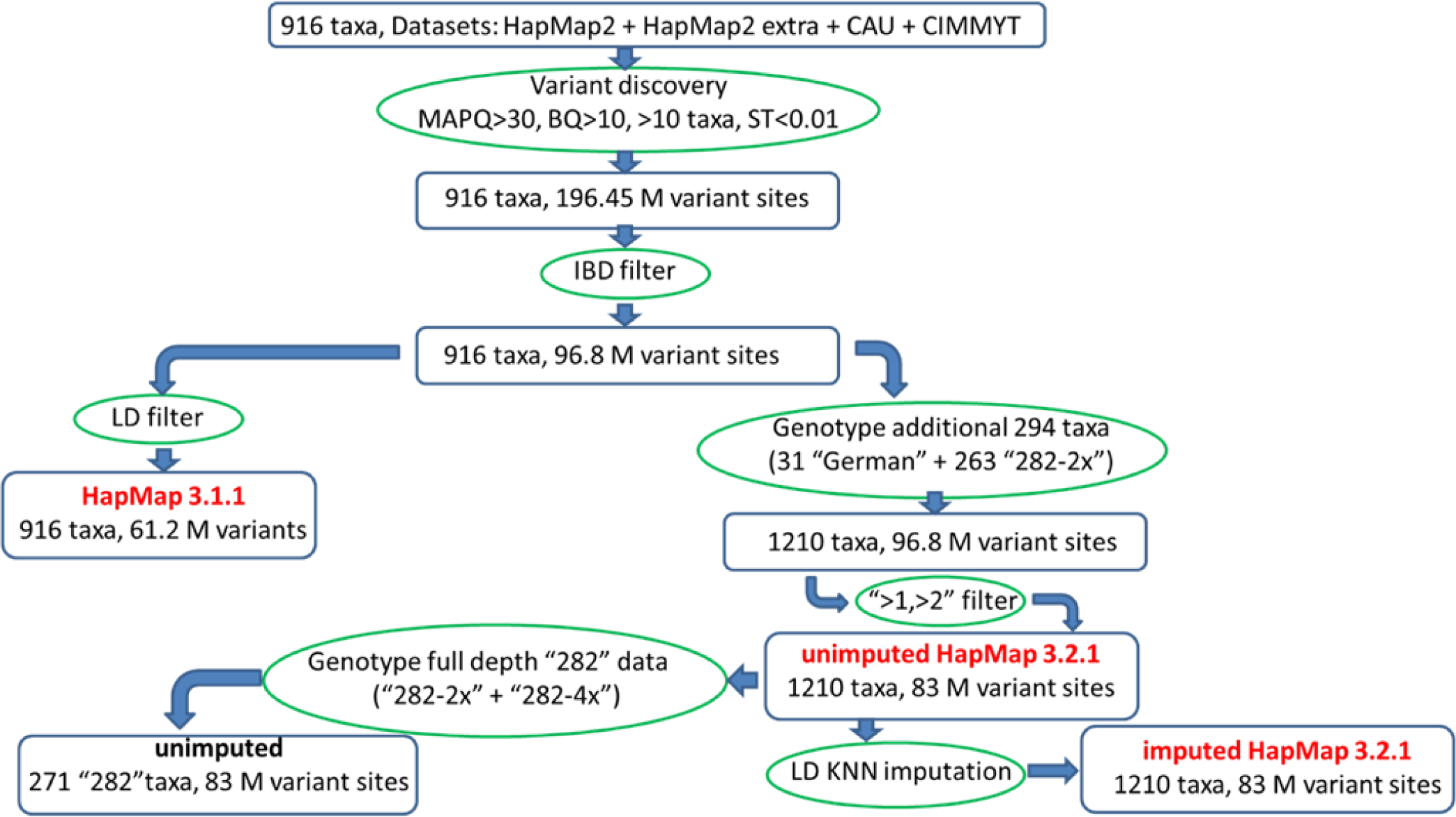
Overview of HapMap 3 pipeline. The exact numbers of variant sites in HapMap 3.1.1 and HapMap 3.2.1 are 61,228,639 and 83,153,144, respectively.

**Table 2:**
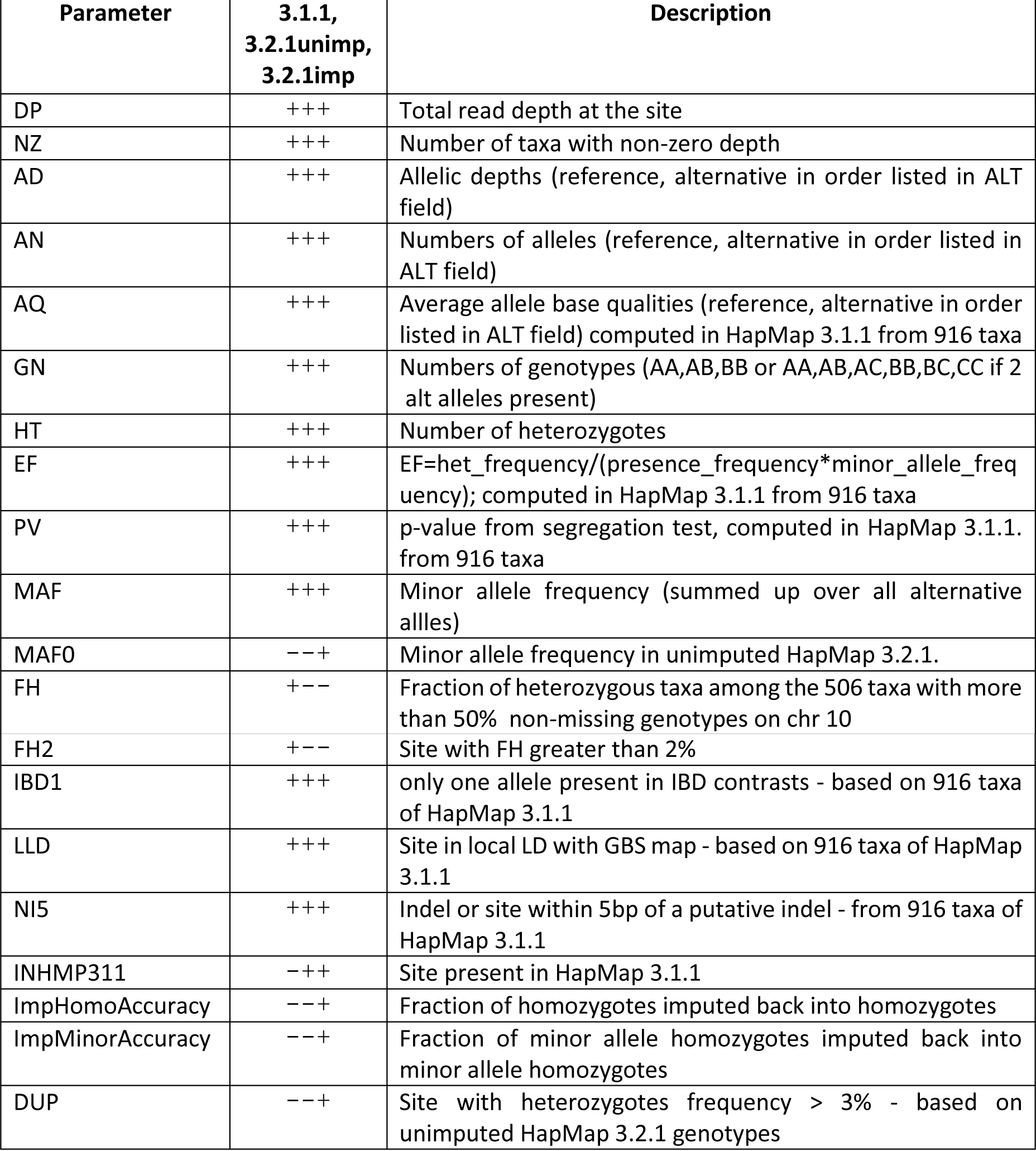
Flags and parameters used in INFO field of VCF files in various HapMap 3 versions

. “+” and “−“ indicate presence or absence, respectively, of a parameter or flag in a given version of HapMap. For example, “-++” means the parameter is present in VCF file of both unimputed and imputed HapMap 3.2.1, and absent from HapMap 3.1.1. VCF files. Unless indicated otherwise, all parameters are computed from depths and genotypes in the current VCF file.

## HapMap 3.1.1

The HapMap 3.1.1 procedure involved checking for linkage disequilibrium of each site against the GBS anchor map [7, 8], which consists of markers located in hypo-methylated and genetically stable regions. Sites giving only very weak or only nonlocal (i.e., outside of 1 Mb radius) linkage Disequilibrium (LD) hits were eliminated, which resulted in the final set of 61,228,639 polymorphisms. For slightly less than 40% of these sites, LD could not be conclusively calculated due to small minor allele frequencies (MAF), whereas the remaining sites, confirmed to be in local LD with the GBS anchor, have been marked with flag “LLD”. Among the sites surviving all filtering steps, 8.7 million are indels or are located near (within 5bp) of an indel. These have been marked with the flag “NI5”. Since a procedure to achieve consistent alignment across all reads covering the same indels - local realignment - has not been performed, genotyping errors could occur, and, consequently, most such sites are tentative and should be treated with caution.

Figure 2 shows overlaps between various classes of variants of HapMap 3.1.1. First, we notice a rather small overlap between sites in confirmed local LD (“LLD” flag) and those marked “IBD1”. This is understandable, as the IBD1 sites represent mostly low MAF cases, where LD assessment could not be done. Indels and vicinity (labeled “NI5”) constitute about 15% of sites in each of the LLD, IBD1, and the union of LLD and IBD1 sets. Only a very small fraction of sites does not carry LLD or IBD1 flag, i.e., they are strongly confirmed by the IBD filter, but could not be classified with LD. The subset of 29.8 million sites in local LD and away from indels should be considered the most reliable.

**Figure 2:**
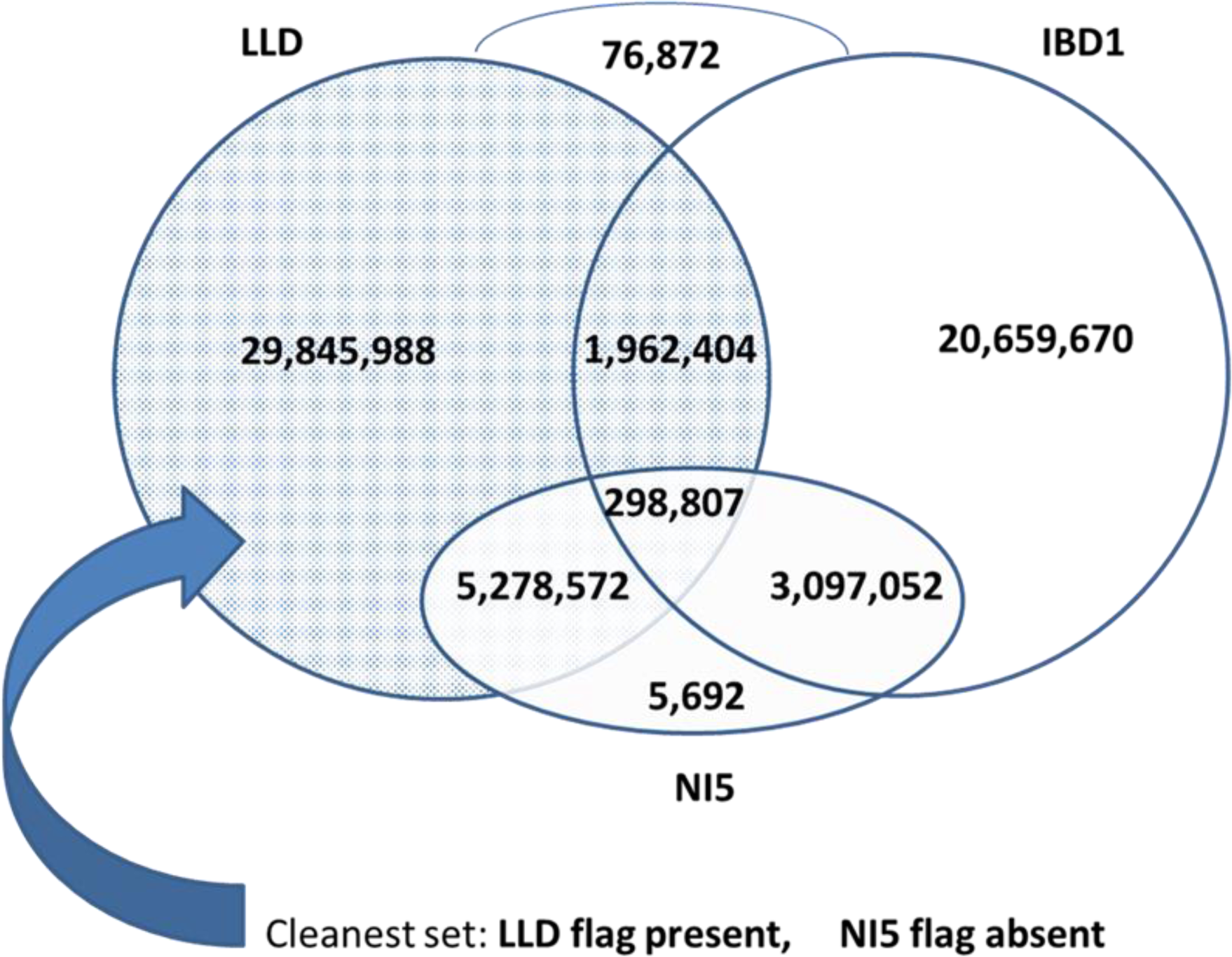
Overlap between various classes of HapMap 3.1.1 polymorphic sites.

To check the sensitivity of the obtained variant set to the mapping quality threshold imposed on the reads counted towards allelic depths, we repeated the pipeline using the mapping quality threshold equal to 1. Comparison of the variant set obtained this way (q1) with our recommended set (q30) is shown in Figure 3. While the overall number of variant sites is approximately independent of the mapping quality threshold, the two pipelines produce significantly different sets of sites, with only 72% of all “q30” sites reproduced by the “q1” pipeline. Closer inspection shows that this variability is due primarily to the IBD1 sites, for which our filtering strategy was the least stringent. On the other hand, the LLD sites, confirmed to be in local LD with GBS anchor, are much more independent of the mapping quality threshold, which confirms high quality of such sites.

**Figure 3:**
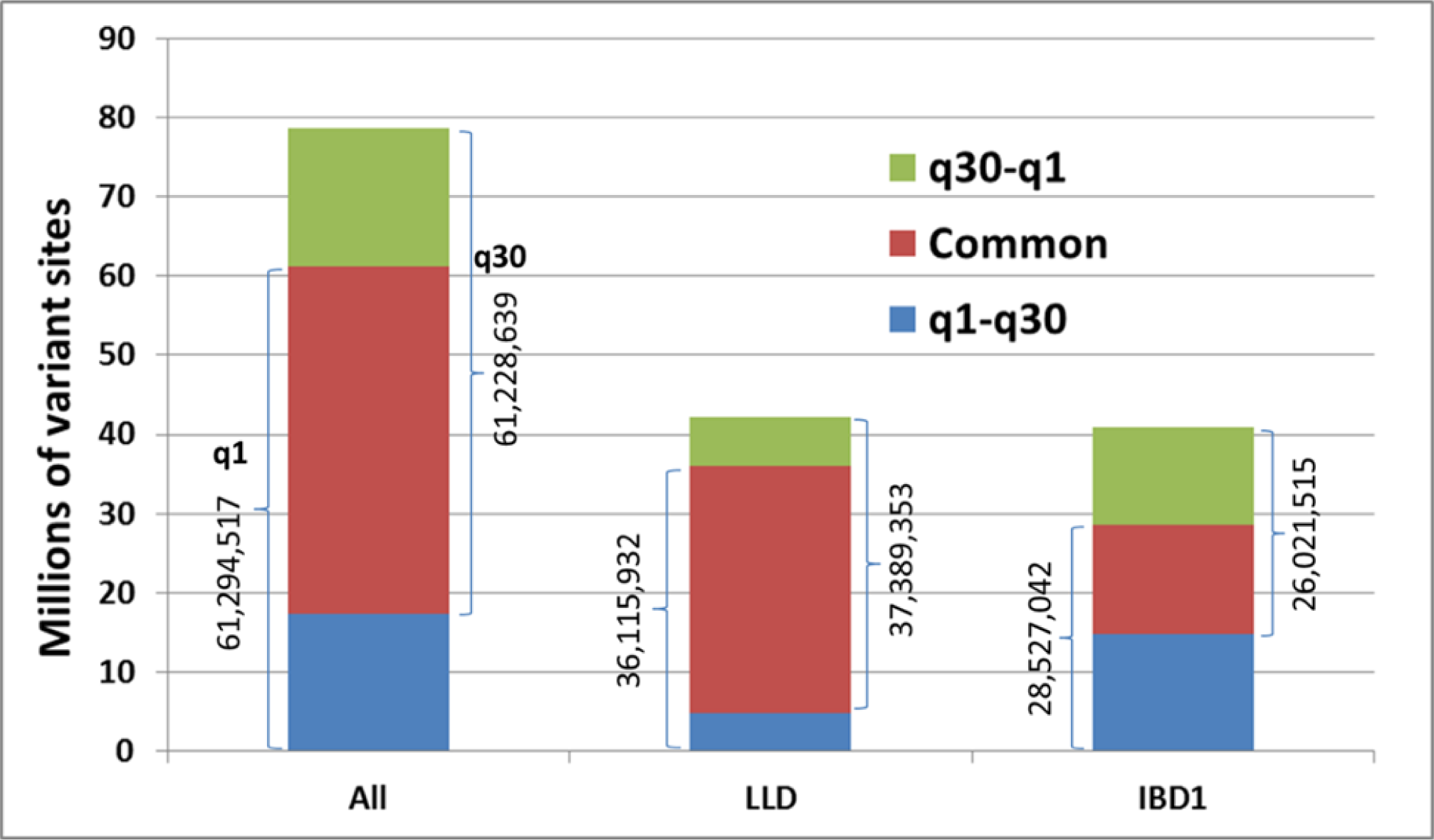
Polymorphic sites detected by HapMap 3 pipeline based on two read mapping quality thresholds.

Importance of choosing a sufficiently tight mapping quality threshold is apparent from Figure 4, where the distribution of inbreeding coefficient computed for chromosome 10 is shown for variant sets obtained with thresholds 1 and 30. The lower threshold results in a large number of miss-mapped reads being counted towards depth, producing overly heterozygous genotypes, especially for highly covered taxa (the peak below 0.8 is due mostly to CIMMYT lines with 10-15x coverage; these lines have higher heterozygosity than other lines which may also contribute to the peak) and thus shifting the curve to the left. Since most HapMap 3 taxa are inbred lines, one should expect the true distribution to be contained within peak around 0.95. In view of this, the “q30” result is definitely an improvement over “q1”, although a longer than expected tail extending towards the value 0.8 indicates that the HapMap 3 variants may contain too many false heterozygotes.

**Figure 4:**
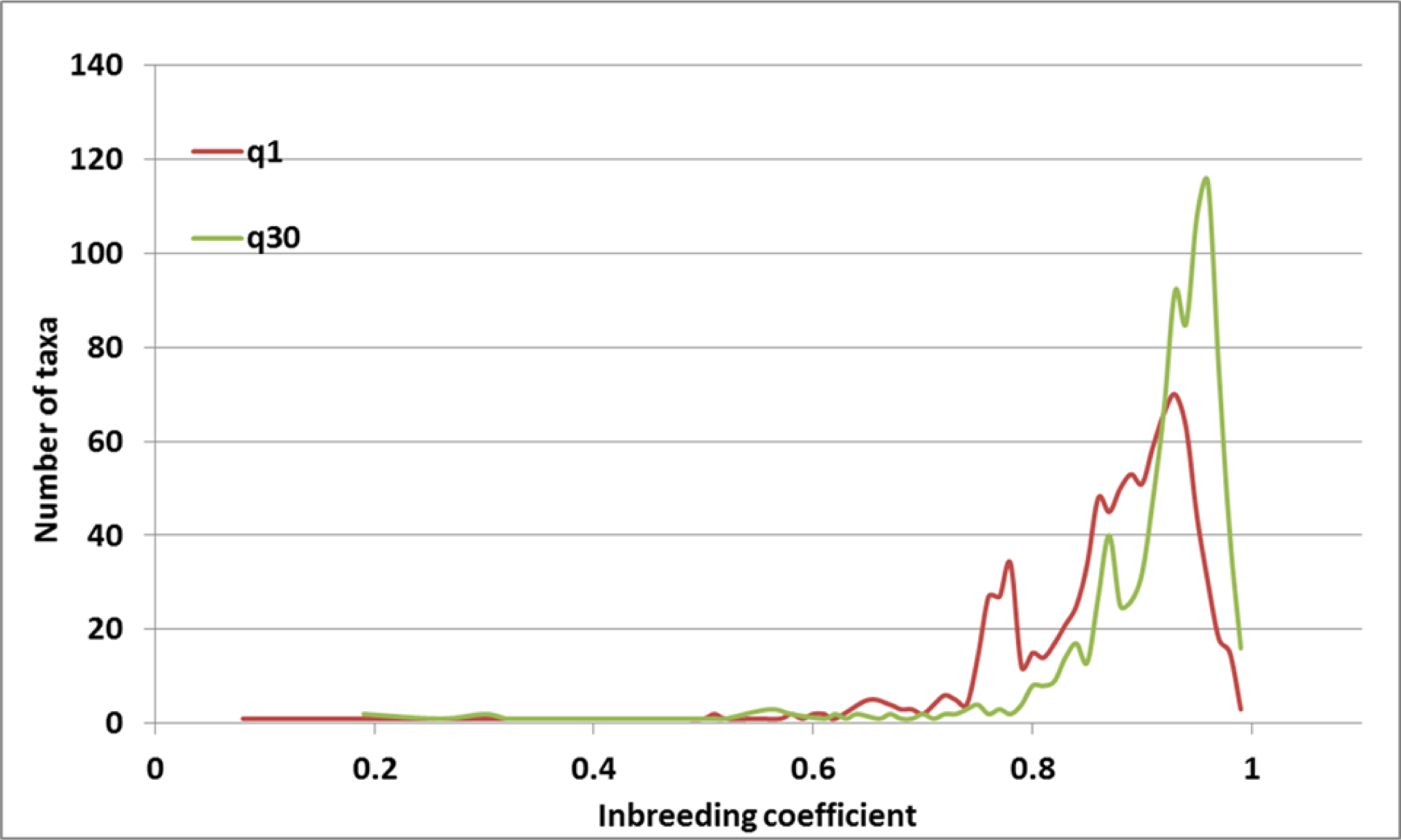
Distribution of inbreeding coefficient for variant sets obtained with two read mapping quality thresholds.

Seemingly heterozygous sites may result from either sequencing errors or misalignments of reads originating from paralogous regions. To investigate this further, we calculated, for each site, the fraction of heterozygous HapMap 3.1.1 genotypes within a subset of 506 high-coverage taxa (defined as those with more than 50% non-missing genotypes on chromosome 10). In HapMap 3.1.1 VCF files, this fraction has been recorded as parameter “FH”. At sites for which this parameter exceeds 2-3%, heterozygotes are likely to originate from misalignments, for example, from tandem and ectopic duplications. Such sites constitute 9% of all HapMap 3.1.1 sites.

## HapMap 3.2.1

The 96.8 million ST- and IBD-filtered variant sites were the starting point for the HapMap 3.2.1 procedure (Figure 1). On these sites, genotypes were called on the 263 taxa from the “282” panel of Ref. [10] using “282 2x” dataset, and on the 31 high-coverage (on average 17 x) “German” taxa [2], for the total of 1210 taxa. Some of the taxa present in the “282” and “German” sets carry the same names as the ones included in the 916-taxa HapMap 3.1.1 set. Since despite identical names such taxa often originate from different germplasm sources, they have been kept separate during genotyping, i.e., reads from different sources were not merged and separate genotypes were computed for each source. In the resulting VCF files, the names of the overlapping taxa have been prefixed by “282set_” and “german_”. For example, in the case of B73, there are three columns representing different datasets for this taxon: “B73” (the original 916-taxa set), “282set_B73” (sequence from the more recent “282” libraries), and “german_B73” (from Ref. [2]).

To further eliminate the false positives resulting from sequencing errors, an additional depth-based filter was applied to the 96.8 million sites. Referred to as “>1,>2” filter it accepts sites for which the read support of minor allele was greater than 1 in at least one taxon and greater than 2 across all taxa. Genotypes on the surviving 83,153,144 sites, referred to as “un-imputed HapMap 3.2.1”, were then processed through the LD KNN imputation procedure based on Ref. [11], where the “nearest neighbors” of a given line are selected based on sites in good local LD with the target site. Whenever possible, the procedure filled up missing genotypes with imputed ones, but the non-missing genotypes were left unchanged, even if imputation classified them differently. Non-imputable missing genotypes at the sites with (pre-imputation) MAF below 1% were assumed to be major allele homozygotes. Imputation reduces the fraction of missing genotypes from 50% to 7%. Most of the originally missing genotypes (about 85%) are imputed to major allele homozygotes. Accuracy of the genotype dataset can be assessed by comparing the original genotypes with imputed ones. As shown in Table 3, 99.8% of major allele homozygotes are imputed back into the same class. While the accuracies of minor allele homozygotes and genotypes including indels are both above 90%, only 11% of heterozygotes are imputed back into the same class, while 47% of them fail imputation altogether. This reflects the inherent difficulty in calling heterozygotes. In the single-reference approach to maize genotyping employed here, heterozygous sites represent true residual heterozygosity as well as misalignments of reads from tandem and ectopic duplications. Since residual heterozygosity in our population of predominantly inbred lines should not exceed 2-3%, all heterozygotes with frequency >=3% can be considered a result of misalignments. About 10% of all heterozygotes present in HapMap 3.2.1 set satisfy this condition. In the VCF files, these sites have been flagged with flag DUP. Other parameters generated by the imputation procedure and recorded for each variant site in the INFO field are ImpHomoAccuracy fraction of all homozygotes imputed back into homozygotes and ImpMinorAccuracy fraction of minor allele homozygotes imputed back to the same class. The INFO field also contains flags IBD1, LLD, and NI5, computed from the initial 916 taxa in the HapMap 3.1.1 procedure. Genotypes resulting from the imputation procedure are referred to as “imputed HapMap 3.2.1”.

**Table 3:** Accuracy of various genotype classes based on statistics from imputation in HapMap 3.2.1

| Genotype class | Accuracy within class [%] | % unimputed |
| --- | --- | --- |
| Major allele homozygote | 99.8 | 1.2 |
| Heterozygote | 11.1 | 47.0 |
| Minor allele homozygote | 94.4 | 14.2 |
| Indel | 92.2 | 17.3 |

Accuracy computed as percentage of the original number of genotypes in a given class (excluding genotypes that could not be imputed) imputed into the same class. The last column shows the fraction of genotypes within a class which could not be imputed.

Relationship between variant sites included in HapMap 3.1.1 and 3.2.1 is shown in Figure 5. Both pipelines start from the same set of IBD-filtered genotypes and subject them to different kinds of filtering, with that of HapMap 3.1.1 being more stringent. It is therefore not surprising that HapMap 3.2.1 recovers the majority (86%) of HapMap 3.1.1 sites, including over 99% of those flagged LLD (i.e., confirmed in local LD). In addition, 30.3 million extra sites are retained in HapMap 3.2.1, which filed the LD filer in HapMap 3.1.1 pipeline. On the other hand, the depth-based “>1,>2” filter applied in HapMap 3.2.1 eliminated 8.2 million sites present in HapMap 3.1.1, including about 0.2 million LLD ones.

**Figure 5:**
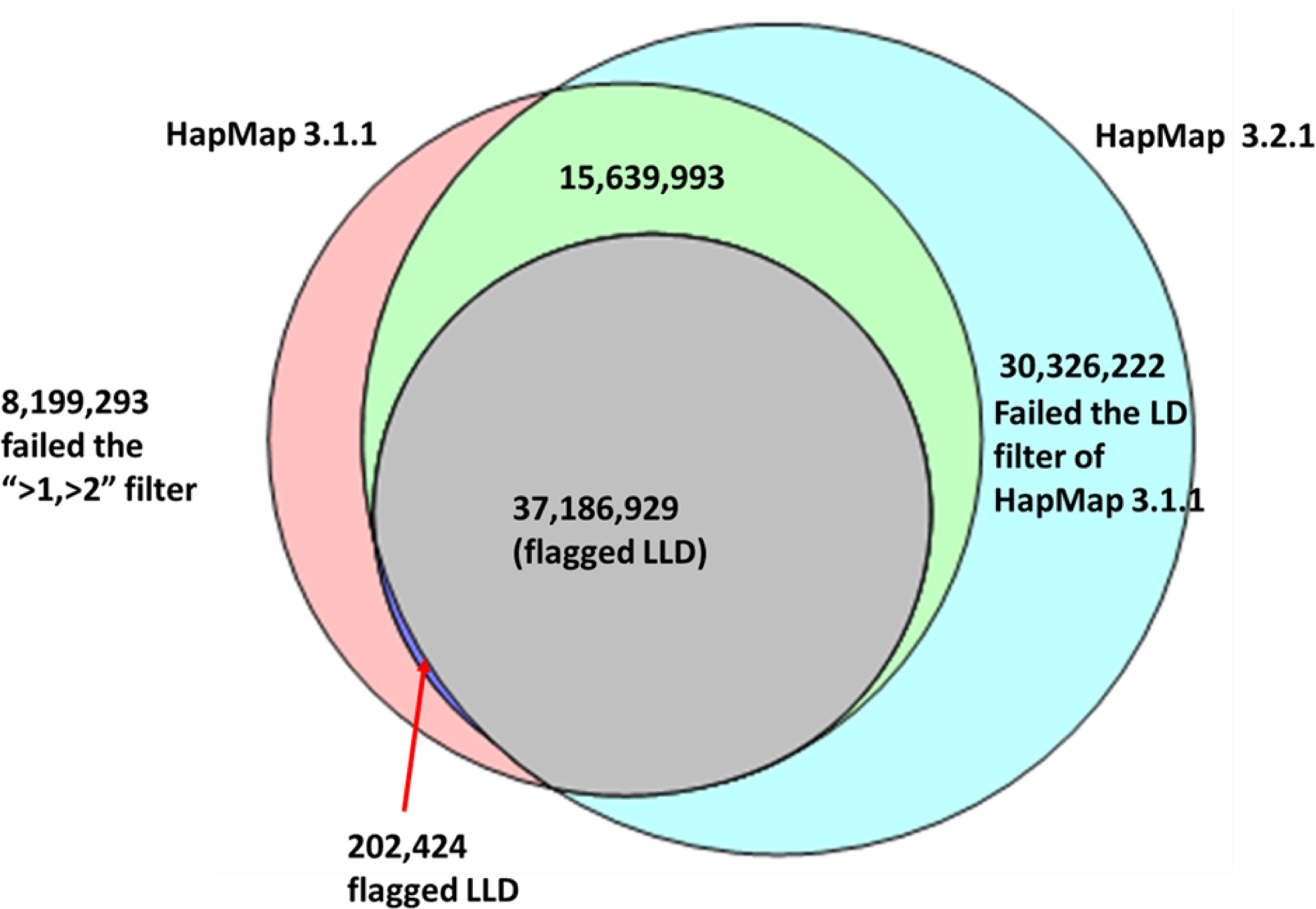
Overlap between HapMap 3.1.1 and HapMap 3.2.1 variant sites.

After the HapMap 3.2.1 release was completed, “282-2x” sequencing data became available for additional 8 taxa from the “282” panel. Libraries for all 271 taxa were also re-sequenced at a higher depth (average of about 4.4x), leading to another dataset, “282-4x” (as this re-sequencing failed for one of the taxa, this dataset only contained 270 taxa). Therefore, the un-imputed HapMap 3.2.1 genotypes for all 271-taxa from the “282” panel were re-called using the full available sequencing depth, creating a separate variant dataset for the “282” panel.

## DISCUSSION

As a species start to diverge, genomic collinearity between individuals deteriorates. The loss of collinearity is by far the biggest challenge to the construction of haplotype maps, because the short sequencing reads used for genotyping do not capture signal for collinearity. The highly repetitive genomic regions are in general easy to identify, because the templates of these repeats are well represented on the reference genome, and sequencing reads mapped to these regions, flagged with low mapping quality, can be removed at the early stage of the analysis pipeline. For HapMap 3, reads with mapping quality lower than 30 were not included in the build. The loss of collinearity of the low copy genomic regions, however, causes vast majority of the genotyping errors, and is not easy to identify computationally, especially for the data sets with limited sequencing depth, which is the case for the maize HapMap 3 project.

The biggest issue in loss of collinearity is the deletion of genomic segment in the individual that is used as the reference genome. The sequencing reads derived from these regions, instead of being removed, would be mapped to other, paralogous regions of the reference genome by the alignment software. In this study, 95%-98% of the reads were mapped by BWA to the reference genome. Many of these mappings are incorrect and result in false positive variant sites. In the human 1000 genome project, a new HaplotypeCaller was used [4], which performs local *de novo* assembly to identify the most likely haplotypes for each individual and thus improve the genotyping results. However, HaplotypeCaller is computationally very expensive, and not always applicable in the species like maize, where reference genome misses much more haplotypes of the pan-genome and has many more paralogous duplications than human. To construct HapMap 3, we relied on the *Zea* GBS map [7, 8], which was obtained from GBS markers located mostly in hypo-methylated chromosomal regions. GBS results were used to identify IBD regions between the individual genomes, and 100 million markers with high percentage of mismatched genotype calls in the IBD regions were filtered out from the initial set of 196 million markers.

The goal of HapMap 3.1.1 is to identify genetic markers in regions where collinearity is preserved in majority of maize lines. The LD filter in the pipeline was applied for this purpose. To do this, we genetically mapped the presence/absence of the minor alleles using the GBS genetic map, and these mapped genetic positions were compared to the physical positions on the B73 reference. Among the 96.8 million sites surviving the IBD filter, 25% did not have enough non-missing data or sufficient minor allele frequency for genetic mapping to be meaningful. For 38% of sites, at least one genetically mapped position matching the physical positons on B73 reference was found, 24% have no significant hits from genetic mapping, probably due to no consensus positions in the HapMap 3 population, and 13% have genetic positions not matching the B73 physical positions. Markers from the latter two categories (37% of all IBD-filtered markers) were removed by the LD filter, leaving slightly over 61 million sites, about 60% of which were confirmed in local LD and marked with a flag “LLD” in VCF files.

The IBD and LD filters applied in the HapMap 3.1.1 project effectively remove majority of the false positive genetic variants caused by paralogous genomic regions, as well as markers with lost collinearity between the species. However, not all the genotyping errors have been removed from the release. 23,839,286 of the sites do not have sufficient minor allele frequency for genetic test (these are missing the “LLD” label in the INFO field of the VCF files). Another source of errors are paralogous regions evolved from tandem duplications. Misalignments of reads from such regions result in false heterozygous genotypes with relatively high frequency and in local LD, and therefore difficult to filter out. Given enough sequencing depth, the tandem duplications can be identified either as copy number variation or imputation errors. However, majority of the HapMap 3 lines have very low sequencing depth, and fail to sample all paralogous loci or all alleles, which makes it difficult to flag all sites complicated by tandem duplications.

Local LD filter based on a large, diverse population may be too stringent, as some markers, good within certain sub-populations, may be thrown out. Therefore, the LD filter was not used in the HapMap 3.2.1 release, which contains a total of 83 million variant sites, subject only to ST and IBD filters and an additional depth-based filter aimed to improve reliability of rare allele calls. Although those sites are likely to have higher misalignment rates, they are still likely to capture real signal related to phenotypic expression.

In the un-imputed HapMap 3.2.1, at about 10% of all variant sites, fraction of heterozygous taxa exceeds 3%. Such sites are marked “DUP”, as most likely originating from duplication misalignments. Figure 6 shows the distribution of fraction of heterozygous sites per taxon for different versions of HapMap 3.2.1 release. While for the un-imputed genotypes the distribution peaks slightly below 1%, imputation significantly shifts the peak to the left, down to about 0.5%. This is a consequence of most missing genotypes being imputed to homozygotes. Interestingly, considering only sites in good local LD (marked with the “LLD” flag) leads to distributions (both imputed and un-imputed) shifted towards higher heterozygosities. This is understandable, as the LLD sites are typically those with higher minor allele frequencies, where the chance of encountering a heterozygote is higher.

**Figure 6:**
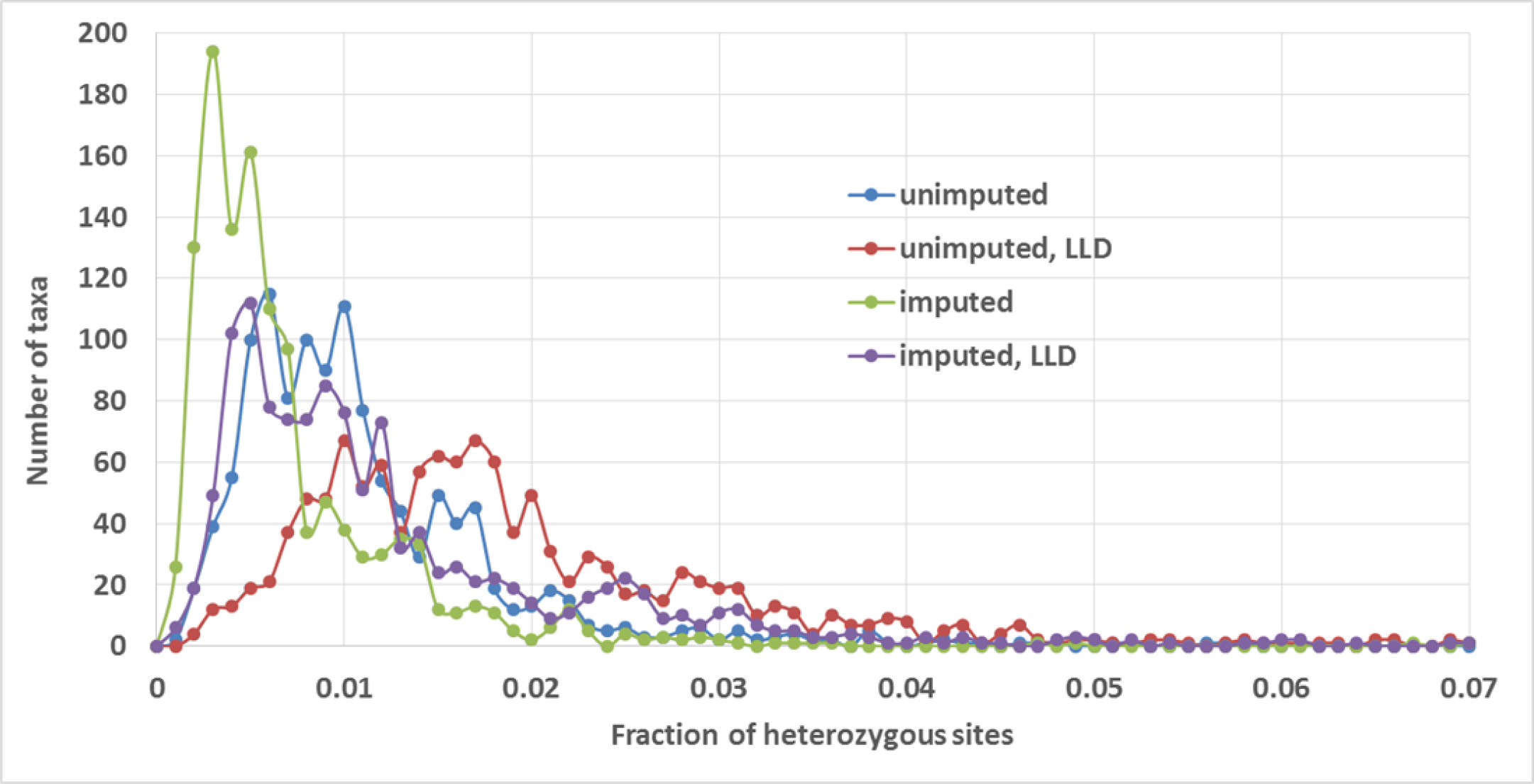
Distribution of fraction of heterozygous sites per taxon for un-imputed and imputed HapMap 3.2.1. Curves marked “LLD” have been obtained considering only sites verified in HapMap 3.1.1 to be in good local LD.

In summary, besides the addition of more maize lines, the HapMap 3.2.1 release differs from the 3.1.1 release in three major aspects: 1) Improved rare allele calls: to increase the accuracy of the variants with rare allele, the HapMap 3.2.1 pipeline applied more stringent read depth thresholds instead of the population genetics based LD filter that could not be applied to sites with very low MAF; 2) The sites with high percentage of heterozygous calls were flagged in the VCF files; 3) Missing data was imputed using the LD KNN method. As summarized in Table 2, the VCF files of both datasets contain labels that flag the characteristics of each of the sites. To effectively use this resource, it is recommended to filter the sites based on the flags that are appropriate to the purpose of each project.

When constructing the maize HapMap 3, the most serious problems we were facing can be attributed to the use of a genome from a single individual (B73) as a reference for other, often very different species. This is becoming the single limiting factor in the study of maize diversity, as well as breeding practice. The only remedy is to move away from a single genome-based reference coordinate and adopt a pan-genome based reference system that incorporates all major haplotypes of the species.

## METHODS

### Plant material

Plant material used in this study was obtained mostly from maize inbred lines representing wide range of *Zea mays* diversity. 103 of these lines, used previously in the HapMap2 project [1], include 60 improved lines, including the parents of the maize nested association mapping (NAM) population [12], 23 maize landraces and 19 wild relatives (teosinte lines, 17 *Z. mays ssp. parviglumis* and 2 *Z. mays ssp. mexicana*). Sequence datasets originating from these lines are referred to in Table 1 as “HapMap2” and “HapMap2 extra”. Majority of the remaining inbred lines originated from CAU (sequence dataset “CAU”) and include, among others, “Chinese NAM” parent lines. Additional 89 inbred lines were provided by CIMMYT and sequenced at BGI (dataset “CIMMYT/BGI”). The HapMap 3 population also contained one *Tripsacum* line (TDD39103), one “mini-maize” line (MM-1A), and a few newly sequenced landraces. Overall, the number of taxa in the initial, variant-discovery stages of the HapMap 3.1.1 project was 916.

The sequence of 271 taxa from the libraries of the “282” panel [10] were added at a later stage (HapMap 3.2.1). DNA to construct these libraries was obtained from the collection that the NCRPIS distributes all over the world. Additionally, the high-coverage data of Ref. [2], originating from 31 European and US inbreds was also included. The total number of taxa genotyped in the HapMap 3.2.1 build is 1218.

During the course of this work several of the taxa present in the sets of germplasm produced by the different members of the consortium carry the same name. Since despite identical names such taxa originated from different germplasm sources, they have all been kept separate for genotyping. For example, it was discovered that new sequence marked as originating from line CML103 actually represents material that is significantly more heterozygous from the line with the same name studied previously in HapMap2 project. Also, the Mo17 sequence originating at CAU has been treated as taxon separate from Mo17 and CAUMo17. In most of those cases, a prefix indicating the origin of the sequence data has been added to the taxa name in order to keep them separated (e.g., “282set_” or “german_”).

## Sequencing

Sequencing has been performed over several years using various generations of Solexa/Illumina instruments and library preparation protocols, giving paired end reads from 44 to 201 bp long. Overall, 113.702 billion reads were obtained, containing 12,497 billion base pairs, giving on average 4.4x coverage per line (assuming 2.3 Gb genome size). However, as shown in Table 1, coverage was not uniform among all lines. For a few lines, sequence generated previously in the context of HapMap2 project was augmented with reads from recent re-sequencing which brought the median coverage of the HapMap2 lines to 5x, with average coverage equal to 7.8x and standard deviation of 7.2x. All NAM parent lines are covered to 10x or better. Most of the 89 lines provided by CIMMYT and sequenced at BGI have coverage exceeding 10x. The recent re-sequencing of the “282” panel resulted in coverage between 1.7 and 36x, averaging 6.5x. Coverage of the 31 “German” lines for Ref. [2] ranges from 8.3 to 59x, with average of 17.4x. Majority of the inbred lines originated from CAU have been sequenced at a lower coverage (1-2x). The list of all lines used in HapMap 3 with the corresponding coverage is given in Additional file 1.

## Alignment

Due to the use of different versions of Solexa/Illumina sequencing equipment, the base qualities in different input FASTQ files are given in different encodings. Prior to alignment, all base qualities have been converted to phred+33 scale. Reads were then aligned to B73 reference (AGP v3) as paired-end using bwa mem aligner (1) with default options. In 72 read sets (Illumina lanes), for technical reasons a high (6%-54%) fraction of paired-end fragments was found to be shorter than reads, so that the two ends contained a part of Illumina adapter and were reverse complements of each other. For such “read-through” fragments, the remnants of Illumina adapter sequences were clipped using TRIMMOMATIC [13] and only one read was used and aligned as single-end. The bwa mem aligner is capable of clipping the ends of reads and splitting each read in an attempt to map its different parts to different location on the reference. As a result, typically over 95% of reads are reported as mapped. However, the fraction of reads with non-zero mapping quality (negative log of the probability that a read has been placed in a wrong location) is much lower – typically only 40-50%. Figure 7 shows a typical distribution of the mapping quality obtained from bwa mem alignment. In practice, we only used alignments with mapping quality of at least 30. A base was counted towards allele depth if its base quality score was at least 10.

**Figure 7:**
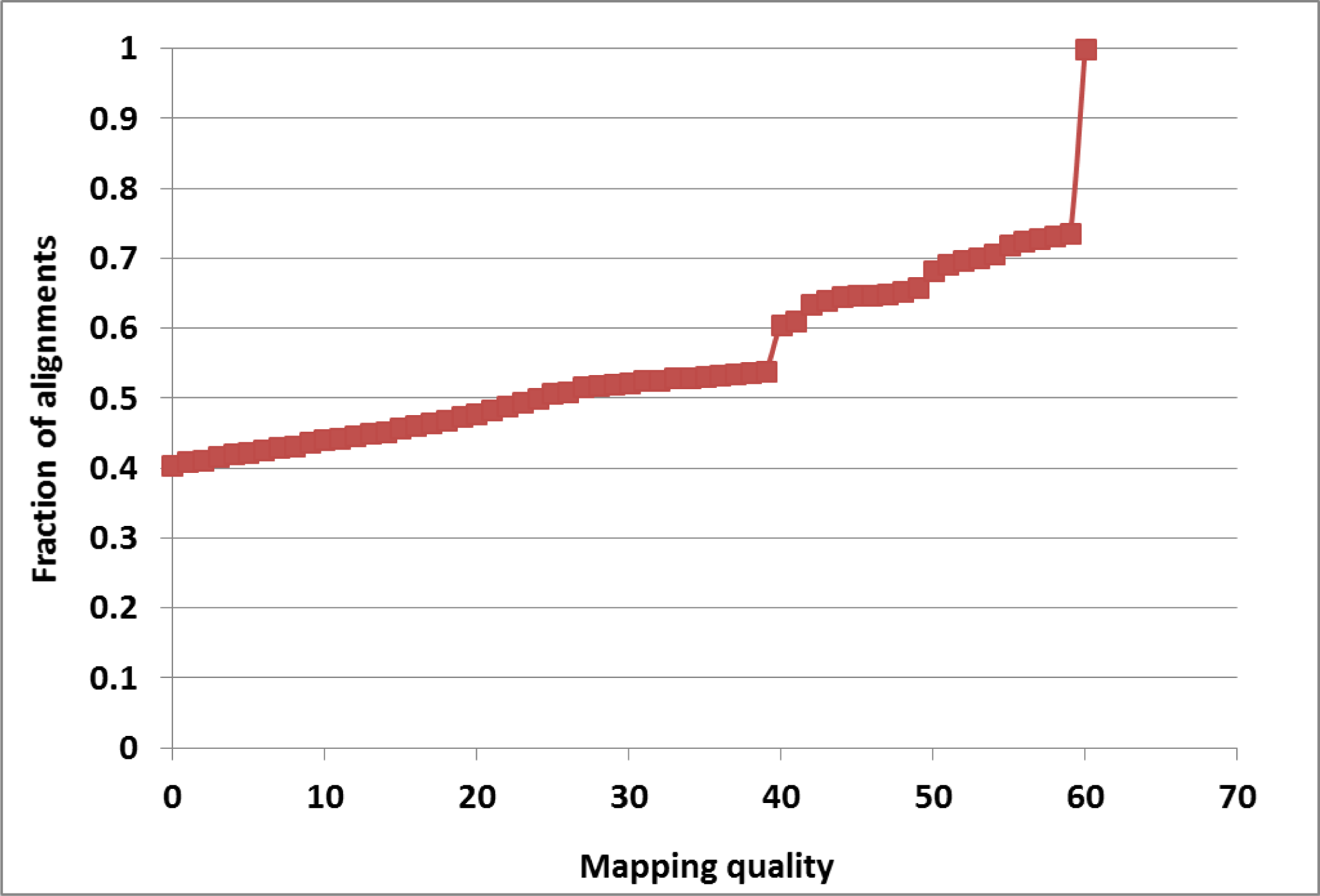
Cumulative distribution of mapping quality from BWA mem alignment of 125,441,950 150bp reads from line A272.

It is well known that alignment may be especially ambiguous when reads contain indels with respect to the reference. In such cases, multiple-sequence realignment approaches have been proposed [4] to find the correct sequence and location of an indel and avoid spurious flanking SNPs. Since indels are not the primary focus of this work and since the realignment is computationally very expensive, it has not been performed by the HapMap 3 pipeline. Thus, although indels and SNPs in their immediate vicinity have been retained in the HapMap 3 VCF files, they are less reliable and have therefore been marked with “NI5” label for easy filtering.

## Genotyping pipeline

Raw genotypes were obtained using a custom-built multi-threaded java code. First, the code executes samtools mpileup command (thresholds on the base and mapping quality are imposed here) for each taxon individually, processing a certain portion of the genome. On a multi-core machine, several such pileup processes (i.e., for several taxa) can be run concurrently as separate threads. Since we are predominantly interested in calling SNPs, we use a simplified indel representation where insertions and deletions with respect to reference are treated as additional alleles “I” and “D”, respectively, regardless of length and actual sequence of the indel. Read depths and average base qualities of all six alleles (A, C, G, T, I, and D) are extracted from samtools mpileup output for each taxon at each genomic position and stored in an array shared between all threads. The amount of memory available on the machine along with the number of taxa determine the upper limit on the size of this array, and therefore – the maximum size of chromosome chunk which can be processed at one time. As base quality of I and D alleles we took the value corresponding to the base directly preceding the indel on the reference.

Extraction of allelic depths for all genomic positions is time consuming, which presents a major obstacle if joint genotyping needs to be re-run, for example, upon extending the taxa set (the so-called “N+1 problem”). It is therefore advantageous to run the depth extraction only once for each taxon and save the obtained depths on disk to be retrieved (rather than re-calculated) during the genotyping step. This way, when the taxa set for genotyping is extended, mpileup step has to be run only for the newly added taxa. Thus, the program features an option to save allelic depths and average qualities in specially designed data structures stored in HDF5 files – one such file per taxon per chromosome. To save space, each allele depth and average quality is stored as one byte, which allows exact representation of integers from 0 to 182, while higher integers (up to about 10,000) are represented approximately by negative byte values through a logarithmic formula with carefully chosen base. Depths and qualities are stored only for sites with non-zero coverage. The details of the storage format and integer representation in terms of byte variables are given in Additional file 2.

Once the allelic depths for all taxa and a given chunk of the genome are available in shared memory, each site is evaluated for presence of a tentative SNP. On a multi-core machine, the set of sites within the genome chunk is divided into subsets processed in parallel on different cores. Sites with less than 10 taxa with read coverage and those with only reference allele present are ignored. For all other sites, genotypes are called for all taxa using a simple likelihood model with a uniform error rate [14] assumed at 1%. Alternative alleles are then sorted according to their allele frequencies and up to two most abundant alleles are kept, as decided by the segregation test described in the next Section. Sites for which all taxa turn out to be reference homozygotes (which may happen despite non-reference alleles being present in the mapped reads) are skipped. Raw variant set obtained in this way is then subject to extensive filtering with the intention of reducing the number of false positives resulting from misalignments.

## Filtering

### Segregation test (ST) filter

For each pair of alleles obtained in the genotyping step, a 2 by N (where N is the number of taxa) contingency table is constructed, containing depths of the first allele in row 1 and depths of the second allele in row 2. The Fisher exact test (FET) is then performed to assess how likely such a table is to occur by chance. If the expected values of the array elements are sufficiently large, the p-value from FET is approximated by that from the computationally efficient chi-square test. However, in most cases encountered here, expensive simulation is needed to obtain sufficiently accurate p-value. To reduce computational burden, we adopted a hybrid approach based on an empirical observation that for statistically insignificant cases (p-values larger than 0.2) the chi-square test results in a de facto lower bound to exact p-values. Thus, the chi-square test is performed first for each site and if the p-value from this test is below 0.2, more exact p-value is obtained from a simulation procedure. The simulation procedure used here, implemented in Java, is the same as the one implemented in R package [15]. An alternative allele is kept if at least one contingency table involving this allele has p-value smaller or equal to 0.01. If none of the alternative alleles survive the ST filter, the site is skipped (not reported in output). The ST filter tends to eliminate variant sites resulting from random sequencing errors.

### GBS anchor map and IBD filter

Given a set of trustworthy SNPs and a diverse set of 916 taxa it is possible to identify, for an arbitrary region of the genome, a number of taxa pairs which are identical by descent (IBD) and are therefore expected to have identical genotypes in this region. If known, these IBD pairs can be used as a powerful filter eliminating variant which violate IBD constraints.

To determine the IBD regions, we used the first step of our pipeline to call genotypes for our 916 taxa on the set of GBS v2.7 sites [7, 8] which tend to concentrate in relatively well-conserved low-copy regions of the genome and can therefore be considered reliable. This set of 954,384 sites was filtered to include only SNP (not indel) sites for which the p-value from the segregation test was below 0.05 and which were more than 5 bp away from any indel. The set of genotypes at 475,272 sites obtained in this way, which will be referred to as GBS anchor, agree well with those from GBS on 167 taxa present in both sets. Alleles detected by the HapMap 3 pipeline agreed with those from GBS at 94% of the GBS sites. At 90% of the sites, fraction of (non-missing data) taxa with genotypes in agreement with those from GBS was at or above 85%. Genotypes different from GBS ones were observed for 82 taxa. These differences were most frequent (up to 19% of all sites) for teosinte lines.

The GBS anchor was used to compute the genetic distance (identity by state) between any two of the 916 lines in windows containing 2000 GBS sites each (about 8.5 Mbp on average). If the genetic distance within such a window was <= 0.02 (about 10 times smaller than the mean distance across all pairs), the two lines were considered to be in IBD. At least 200 comparable GBS sites (i.e., non-missing data simultaneously on both lines being compared) were assumed necessary to make the genetic distance calculation feasible.

The number of taxa involved in IBD relationships in any given window were between 385 (start of chromosome 10) and 757 (middle of chromosome 7) and averaged 588, leading to large numbers of IBD contrasts, ranging from 3,710 (beginning of chromosome 4) to 42,890 (middle of chromosome 7), and averaging 13,500.

The raw (ST-filtered) genotypes were checked against the IBD pairs in various regions, using a procedure which counts, for each site, numbers of base matches and mismatches for each allele present at the site. If the match/mismatch ratio is at least 2 for at least two alleles, or if only one allele is present in all IBD contrasts, the site is considered as passing the IBD filter. Such a filter is less powerful for sites where all bases in IBD lines are major allele homozygotes (i.e., the SNP being evaluated occurs in lines not involved in IBD pairs). Formally, such a site passes IBD filter, but this is statistically easier to achieve than agreement involving minor alleles, so the actual SNP is not strongly confirmed. These uncertain sites, mostly with low minor allele frequency, are labeled “IBD1” in the HapMap 3 VCF files and constitute about 50% of all HapMap 3 sites.

### Linkage Disequilibrium (LD) filter

Any true SNP should be in local linkage with other nearby SNPs. This observation gives rise to another filter used in this work, referred to as the LD filter. For each variable site surviving the ST and IBD filters, we evaluate LD with each site of the GBS anchor. As the LD measure we chose the p-value from a 2 by 2 contingency table of taxa counts corresponding to the four haplotypes (AB,Ab,aB,ab). For simplicity, heterozygous genotypes were treated as homozygous in minor allele. For a pair of sites to be tested for LD, the following three conditions had to be satisfied to make the calculation meaningful: i) the two sites were at least 2,500bp apart, ii) there were at least 40 taxa with non-missing genotypes at both sites being compared, and iii) at least 2 taxa with minor allele had to be present at each of the two sites.

Filtering procedure executed for each site is summarized in Figure 8. First, LD between the given site and all sites in GBS anchor was computed and up to 20 best LD hits (the ones with lowest p-values) were collected. If the p-value of the best hit exceeded 1E-6 (which roughly corresponds to the peak of the overall distribution of p-values), the site was rejected. Otherwise, it was determined whether the set of best hits contained any local hits, i.e., hits to GBS sites on the same chromosome within 1 Mbp of the site in question and with the p-value smaller than 10 times the p-value of the best hit. If no such local hits were found, the site was rejected, otherwise it was kept and marked as a site in Local LD using the flag “LLD”. Note that the procedure defined this way filters out sites with only non-local LD hits as well as those with only weak LD signal. Sites in local LD as well as those for which LD could not be assessed (because of low minor allele frequency or missing data) pass the filter.

**Figure 8:**
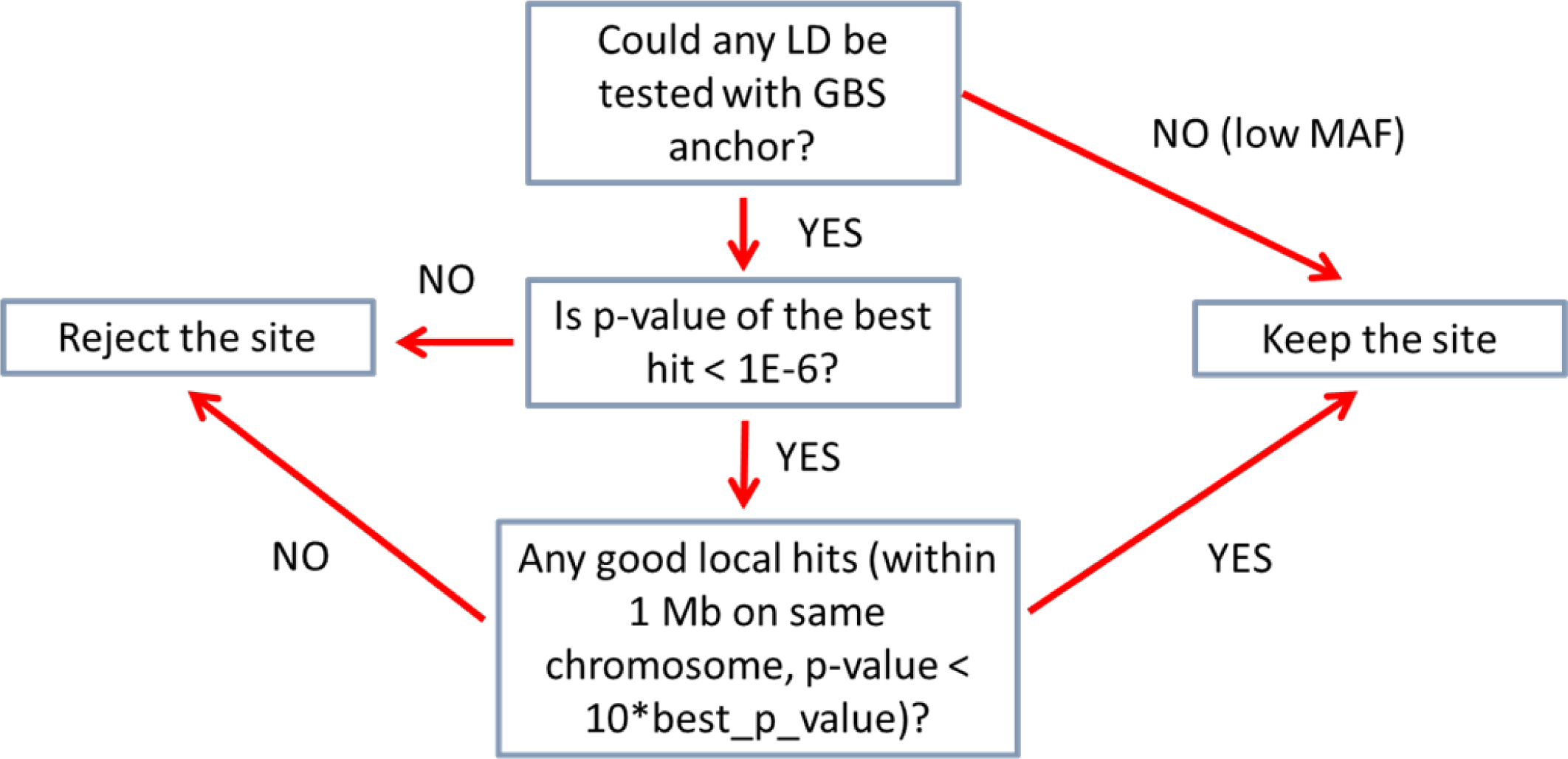
Linkage Disequilibrium-based filtering flowchart.

## Imputation

In the HapMap 3.2.1 pipeline, the ST- and IBD-filtered genotypes, after the application of the additional “>1,>2” depth-based filter, were processed through the LD KNN imputation procedure based on Ref. [11] to fill in the missing data. The procedure is a version of the “K nearest neighbors” routine where the “nearest neighbors” of a given taxon are selected based on genetic distance computed using variant sites in good local LD. Specifically, for a given target site, a list of up to 70 sites in best LD (as given by the *R*^2^ measure) with it is compiled by checking all surrounding sites within 600Kb characterized by heterozygosity lower than 3% and more than 50% taxa with non-missing genotypes. Then, at the same target site, for each target taxon, up to 30 “nearest neighbor” taxa are selected, giving the lowest genetic distance from the target taxon. Genetic distances are computed using the set of local LD sites selected in the previous step. Taxa with less than 50% non-missing genotypes at LD sites, missing genotype at the target position, having distance from the current taxon larger than 0.1, or resulting in less than 10 common LD sites on which the distance can be calculated, are excluded from distance calculation process. Genotypes of the selected nearest neighbor taxa at the target site are stored in memory along with the genetic distances from the target taxon. This information is the used to compute a weight *W*_*i*_ of each neighbor genotype *g* as follows:

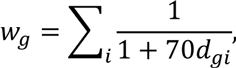

where the summation index *i* runs over all neighboring taxa with genotype *g* at the target site, and *d*_*gi*_ is the distance of taxon *i* from the target taxon. The genotype with the highest weight is considered the imputed genotype (of the target taxon at the target site) provided its weight is at least 10 times larger than that of the second-best candidate genotype. Otherwise the imputation is considered inconclusive and the imputed genotype is set to “unknown” (missing data), as it is in the case when no close neighbors of the current taxon could be found. If a genotype imputed to “unknown” occurs at a site where MAF<1%, it is automatically converted into major allele homozygote.

The imputation procedure is run for each genotype in the input file. However, in the output only the originally missing genotypes are updated to imputed ones, whereas all others are left unchanged, even if classified differently. On the other hand, all imputed genotypes are used during a run to collect imputation statistics. The “transition matrix” showing how many genotypes originally in a given class were imputed into other classes is an indication of the accuracy of the input genotypes. Error rates calculated from the transition data are given in Table 3.

## AVAILABILITY OF DATA

At present, reads from all datasets is available in the form of BAM files (with reads aligned to AGP v3 reference) on CYVERSE data store (formerly iPlant, http://www.cyverse.org/data-store), in directories `/iplant/home/shared/panzea/raw_seq_282/bam` (dataset “282-2x” and “282-4x”), `/iplant/home/shared/panzea/hapmap3/bam_germanlines` (“German” dataset), and `/iplant/home/shared/panzea/hapmap3/bam` (other datasets). Raw reads from the “German” dataset are also available from NCBI BioProject with accession PRJNA260788.

The set of HapMap 3.1.1. polymorphisms determined for 916 taxa (from datasets “HapMap2”, “HapMap2 extra”, “CAU”, and “CIMMYT/BGI”) is available in VCF format on CYVERSE data store in the directory `/iplant/home/shared/panzea/hapmap3/hmp311`, in files `c*_hmp311_q30.vcf.gz` (one file per chromosome, where “*” stands for chromosome 1-10). Additionally, files `c*_hmp311_q1.vcf.gz` (in the same location) contain test results obtained with mapping quality threshold equal to 1.

The HapMap 3.2.1 variants for 1210 taxa (916 initial Hapmap 3.1.1 taxa + 263 taxa from “282-2x” set + 31 “German” lines) are available from `/iplant/home/shared/panzea/hapmap3/hmp321/unimputed`, in files `merged_flt_c*.vcf.gz` (un-imputed results) and in `/iplant/home/shared/panzea/hapmap3/hmp321/imputed`, in files `merged_flt_c*.imputed.vcf.gz` (imputed results). Files `c*_282_onHmp321.vcf.gz` in CYVERSE directory `/iplant/home/shared/panzea/hapmap3/hmp321/unimputed/282_libs_2015` contain un-imputed genotypes on HapMap 3.2.1 sites from the full depth data available for the “282” panel (271 taxa, datasets “282-2x” + “282-4x”).

## LIST OF ABREVIATIONS

International Maize and Wheat Improvement Center (CIMMYT), Segregation test (ST), Identity-by-descent (IBD), Genotyping-by-sequencing (GBS), Linkage Disequilibrium (LD), Minor Allele Frequency (MAF), Nested Association Mapping (NAM), Single Nucleotide Polymorphism (SNP), Insertion-Deletion (Indel)

## ADDITIONAL FILES

Additional file 1: HapMap3TaxaAndCoverage.xlsx – spreadsheet with a list of all lines used in HapMap 3 with their corresponding coverage

Additional file 2: DepthFormatDetails.pdf - details of byte representation and storage format used for allelic depths

